# Perfusion-Independent Tissue Hypoxia in Cardiac Hypertrophy in Mice Measured by ^64^Cu-CTS PET Imaging

**DOI:** 10.1101/2024.04.22.590587

**Authors:** Friedrich Baark, Aidan M. Michaels, Edward C. T. Waters, Alex Rigby, Jana Kim, Zilin Yu, Victoria R. Pell, James E. Clark, Philip J. Blower, Thomas R. Eykyn, Richard Southworth

## Abstract

**Background:** Hypoxia is central to many cardiac pathologies, but clinically its presence can only be inferred by indirect biomarkers including hypoperfusion and energetic compromise. Imaging hypoxia directly could offer new opportunities for the diagnosis and sub-stratification of cardiovascular diseases.

**Objectives:** To determine whether [^64^Cu]CuCTS Positron Emission Tomography (PET) can identify hypoxia in a murine model of cardiac hypertrophy.

**Methods:** Male C57BL/6 mice underwent abdominal aortic constriction (AAC) to induce cardiac hypertrophy, quantified by echocardiography over 4 weeks. Hypoxia and perfusion were quantified in vivo using [^64^Cu]CuCTS and [^64^Cu]CuGTSM PET, respectively, and radiotracer biodistribution was quantified post-mortem. Cardiac radiotracer retention was correlated with contractile function (measured by echocardiography), cardiac hypertrophy (measured by histology), HIF-1α stabilization and NMR-based metabolomics. The effect of anesthesia on [^64^Cu]CuCTS uptake was additionally investigated in a parallel cohort of mice injected with radiotracer while conscious.

**Results:** Hearts showed increased LV wall thickness, reduced ejection fraction and fractional shortening following AAC. [^64^Cu]CuCTS retention was 317% higher in hypertrophic myocardium (p<0.001), despite there being no difference in perfusion measured by ^64^CuGTSM. Radiotracer retention correlated on an animal-by-animal basis with severity of hypertrophy, contractile dysfunction, HIF1α stabilization and metabolic signatures of hypoxia. [^64^Cu]CuCTS uptake in hypertrophic hearts was significantly higher when administered to conscious animals.

**Conclusions:** [^64^Cu]CuCTS PET can quantify cardiac hypoxia in hypertrophic myocardium, independent of perfusion, suggesting the hypoxia is caused by increased oxygen diffusion distances at the subcellular level. Alleviation of cardiac workload by anesthesia in preclinical models partially alleviates this effect.

## Introduction

Hypoxia is a central component of cardiac ischemia. It is an important factor in coronary microvascular disease, cardiac hypertrophy (1) and heart failure (2), as well as being a key driver of angiogenesis (3). It is thought to underlie and precede many biochemical, structural and functional changes during chronic cardiovascular diseases (4). While the presence of tissue hypoxia in cardiovascular disease has long been inferred using a variety of proxies including loss of perfusion, stabilization of HIF1α and measurement of metabolic derangement via numerous means(5), it is perhaps surprising that intracellular hypoxia has only ever successfully been directly measured in one cardiac patient, to our knowledge (6). Ischemia and hypoxia are defined as a mismatch between supply and demand of perfusion and oxygenation, respectively (7). Quantification of cardiac perfusion/coronary flow reserve by magnetic resonance or nuclear imaging techniques are well established but provide only one half of the equation i.e. supply. With no reference to “demand” in the subtended tissues, the pathophysiological relevance of measured hypoperfusion is difficult to interpret. Similarly, stabilization of HIF1α and cardiac metabolic derangement are indicative of hypoxia but not specific to it (8,9), and in any case are not easily obtainable biomarkers clinically.

Non-invasive imaging of hypoxia itself could provide new opportunities for the diagnosis and sub-stratification of cardiovascular disease by quantifying the *mismatch* between oxygen supply and demand. This may be more selective than quantifying perfusion alone, particularly for cardiovascular diseases associated with microvascular dysfunction where ischaemia is patchy, diffuse, and hard to detect by any other means (10,11). Copper bis(thiosemicarbazones) (BTSCs), radiolabeled with either ^64^Cu or ^62^Cu, show promise as PET hypoxia imaging agents. They have been widely investigated for cancer imaging (12), but remain relatively underexplored in cardiology. The lead BTSC compound [^64^Cu[CuATSM (^64^Cu-2,3-butanedione bis(N4-methylthiosemicarbazone)) has been shown to deposit ^64^Cu in hypoxic cardiac myocytes (13), isolated perfused hearts (14), and in regionally occluded canine myocardium *in vivo* (15). However, in the one small clinical trial (7 patients) that has been performed to date, [^62^Cu]CuATSM was only able to delineate cardiac hypoxia in one patient with unstable angina, despite 4 of those patients exhibiting elevated ^18^FDG uptake (16). Thus, while [^64/62^Cu]CuATSM PET can detect severe degrees of hypoxia in tumors and extreme experimental models of cardiac ischaemia, it is not sensitive enough to detect the subtle hypoxia thought to characterize chronic cardiac ischemic syndromes.

In recent years we have screened and validated a library of BTSC complexes, some of which may be more sensitive to the less extreme degrees of hypoxia relevant to cardiology. Of these, [^64^Cu]CuCTS ([^64^Cu]copper-2,3-pentanedione-bis(thiosemicarbazone) complex) exhibits a hypoxic:normoxic retention ratio of 16:1 (compared to 8:1 for [^64^Cu]CuATSM), and crucially, deposits radiocopper in isolated perfused rat hearts where they start to exhibit loss of energetic homeostasis (by ^31^P MR spectroscopy), elevated lactate washout (indicating low-level anaerobic glycolysis), HIF1α stabilization and hypocontractility at the “tipping point” which [^64^Cu]CuATSM cannot identify (17).

In this study, we determine whether [^64^Cu]CuCTS can be used to image hypoxia *in vivo* in a pathophysiologically relevant rodent model of chronic cardiovascular hypertrophy. We hypothesize that non-compensated hypertrophic myocardium is hypoxic (i.e. oxygen-deprived due to capillary rarefaction and increased diffusion distances across larger myocytes), rather than being due to loss of perfusion. If it can, this imaging approach could offer new diagnostic opportunities for hard-to-characterize coronary microvascular diseases and offer new insight into cardiovascular disease more widely, particularly those currently classified as “non-ischemic”, despite only perfusion or coronary flow reserve having been measured to date.

## Methods

### [^64^Cu]CuCTS Radiolabeling

^64^Cu was produced at the University of Birmingham, UK on an MC-40 cyclotron and purified at KCL as previously described. 2,3-pentanedione-bis(thiosemicarbazone) (CTS) was synthesised in-house and radiolabeled as previously described (18), achieving >98% radiochemical purity, confirmed by thin layer chromatography.

### Animals

Male C57BL/6 mice (22-25g, Charles River, UK) were used for all experiments. Procedures were approved by KCL’s local Animal Care and Ethics Committee and carried out in accordance with Home Office regulations as detailed in the Guidance on the Operation of Animals (Scientific Procedures) Act 1986.

### Echocardiography

Longitudinal monitoring of cardiac function was performed using a Vevo 3100 echocardiography system (VisualSonics, Toronto, Canada) with an MX400 transducer at 30 MHz. Mice were anesthetized and maintained using isoflurane (1.5-2.0 % in O_2_) at 37° on a homeothermic platform. Parasternal left ventricle (LV) long-axis and short-axis M-mode and B-mode images were analyzed using Vevo software (VisualSonics) to calculate cardiac function, mass and geometries. M-mode images were captured at the level of the papillary muscle.

### Cardiac hypertrophy induction by abdominal aortic constriction (AAC)

Mice were anaesthetized with isoflurane (1.5-2.0 % in O_2_) on a heating pad to maintain body temperature at 37°C, and the chest was shaved and disinfected with Videne. The abdominal aorta was exposed by laparotomy, a blunted 27G needle placed on top, and 8-0 suture (W2808, Ethicon) tied around aorta and needle. The needle was then withdrawn, leaving the looped suture restricting the aorta (19). The abdominal wall and skin were closed with 6-0 vicryl sutures and the surgical site cleaned with Videne. Mice recovered in a heated chamber with buprenorphine (20 µg/kg, Vetergesic, Ceva Animal Health Ltd, France) for perioperative analgesia. Sham animals underwent the same procedure, but the aorta was not tied.

### Evaluation of [^64^Cu]CuCTS PET for imaging hypoxia in AAC-induced cardiac hypertrophy

Mice were anaesthetized with isoflurane (1.5-2% in O_2_) on a heated platform and injected via tail vein with [^64^Cu]CuCTS (100 ± 50 µL, 1.0-3.5 MBq), and radiotracer allowed to circulate for 20 minutes. Mice were scanned in a nanoPET/CT scanner (Mediso, Hungary) using a four-bed “mouse hotel” system. CT (480 projections; helical acquisition; 55 kVp) was performed for anatomical visualization. Thirty minutes later a PET acquisition (2 x 15 min 1:5 coincidence mode) was performed. Radiotracer concentration was quantified using VivoQuant v 2.5 (InVicro Ltd.) using CT to define the ROI. A parallel cohort (n = 4 plus sham) were subjected to the same protocol, but imaged with the perfusion tracer [^64^Cu]CuGTSM (glyoxalbis(N(4)-methyl-3-thiosemicarbazonato) copper(II)) for comparison. Mice were then culled and organs excised for *ex vivo* radiotracer γ-counting.

### Investigation into the effect of anesthesia on cardiac ^64^Cu deposition

The use of anesthesia lowers cardiac workload, and could therefore lead to an underestimation of the sensitivity of [^64^Cu]CuCTS to hypoxia. To investigate this, a further group were immobilized in a mouse restrainer (Braintree Scientific) to allow [^64^Cu]CuCTS injection without anesthesia, and returned to their cage conscious for 20 min prior to PET imaging under isoflurane. Cardiac [^64^Cu]CuCTS retention was compared to a parallel group anesthetized with isoflurane throughout as a time-matched control.

### Immunohistochemistry

Immunohistochemistry to assess myocyte size (by wheatgerm agglutinin), β-myosin heavy chain expression and HIF1α nuclear staining was performed within 7 days of tissue harvesting (see Supplementary Information). Sections were imaged using a Leica TCS SP5 II confocal microscope and analysed using LAS X Life Science Software.

### HIF1α quantification by Western blotting

Tissues were crushed to powder under liquid nitrogen and processed as detailed in the Supplementary Information. Blots were probed for HIF1α (NB100, 1:500; Novus Biologicals) with β-actin as a loading control with HRP-linked anti-rabbit IgG secondary antibody (1:10,000; Abcam). Densitometry was performed using Quantity One Software (BioRad), normalized to β-actin.

### Quantification of blood troponin and brain natriuretic peptide levels

Mouse cardiac troponin I (CTNI-1-HS, Life Diagnostics) and mouse BNP (NBP2-70011, Novus Biologicals) levels in serum were quantified using ELISA as per manufacturers’ instructions.

### NMR metabolomics

Metabolites were extracted from frozen tissue as previously described (20). Tissue was homogenized in ice-cold methanol, and ddH_2_O and chloroform were added in a 1:1:1 ratio. Samples were centrifuged for 1 h at 2500g at 4 °C, the resultant aqueous upper layer was aspirated, 20-30 mg chelex-100 was added to chelate paramagnetic ions and the homogenate was vortexed and centrifuged at 2500g for 5 min at 4°C. Supernatants were mixed with 20 µL universal indicator prior to freeze-drying overnight. The freeze-dried pellet was then reconstituted in 600 μL of deuterium oxide [containing 8 gL^−1^ NaCl, 0.2 gL^−1^ KCl, 1.15 gL^−1^ Na_2_HPO_4_, 0.2 gL^−1^ KH_2_PO_4_ and 0.0075% w/v 3-(trimethylsilyl)-2,2,3,3-d_4_-tetradeuteropropionic acid (TSP)] and adjusted to pH ~7 by titrating with 100 mM hydrochloric acid. NMR acquisition parameters are given in the Supplemental Information.

### Statistics

Analysis was performed blinded and randomized using GraphPad Prism 9. All data are presented as mean ± standard deviation unless otherwise stated. Comparisons between results were analyzed using unpaired t-tests (with Welch’s correction) and one-way or two-way ANOVA with Tukey correction. Results were considered statistically significant with a *p*-value <0.05. Unclassified principal components analysis (PCA) was performed in Matlab. Metabolite peak integrals were first normalized to total spectrum area prior to PCA calculation using Venetian blinds cross validation and five cross validation groups. Hierarchical cluster analysis was performed in Matlab using the function clustergram.

## Results

### Progression of cardiac hypertrophy after abdominal aortic constriction

An initial longitudinal study was performed to identify the most suitable timepoint for evaluating PET tracer performance during the progression of hypertrophy. Mice (n = 6/group) were randomly assigned into AAC and sham groups. Animals were monitored by echocardiography weekly, and cohorts (with sham controls) were sequentially culled 1, 2, 4, 6 and 10 weeks after surgery. Their hearts were excised and immediately fixed and frozen in OCT for histology/immunohistochemistry/Western blot analysis for biomarkers of hypoxia. AAC surgery caused cardiac hypertrophy which peaked at week 6 before progressing to ventricular dilatation over the following 6-10 weeks, associated with progressive increases in myocyte cross-sectional area and β-myosin staining which were significantly elevated from 1-week post-surgery **(see data supplement)**. Cardiac HIF1α stabilization and serum BNP concentrations were biphasic; both were evident 1 week after surgery, with serum BNP peaking at 2 weeks and cardiac HIF1α peaking at 4 weeks. Based on this experiment, 4 weeks post-surgery was selected as the optimal timepoint to evaluate the sensitivity and selectivity of [^64^Cu]CuCTS to cardiac hypoxia in the subsequent study.

After baseline hemodynamics were established by echocardiography, 12 mice underwent AAC surgery alongside 9 sham controls. Four weeks later, AAC-operated mice displayed significantly elevated cardiac mass, loss of ejection fraction and fractional shortening, increased septal and left ventricular free wall thickness (Figure 1) consistent with our prior range-finding study, associated with significantly increased cardiomyocyte cross-sectional area, β-myosin heavy chain expression and elevated nuclear staining for HIF1α (Figure 2).

**Figure 1.**
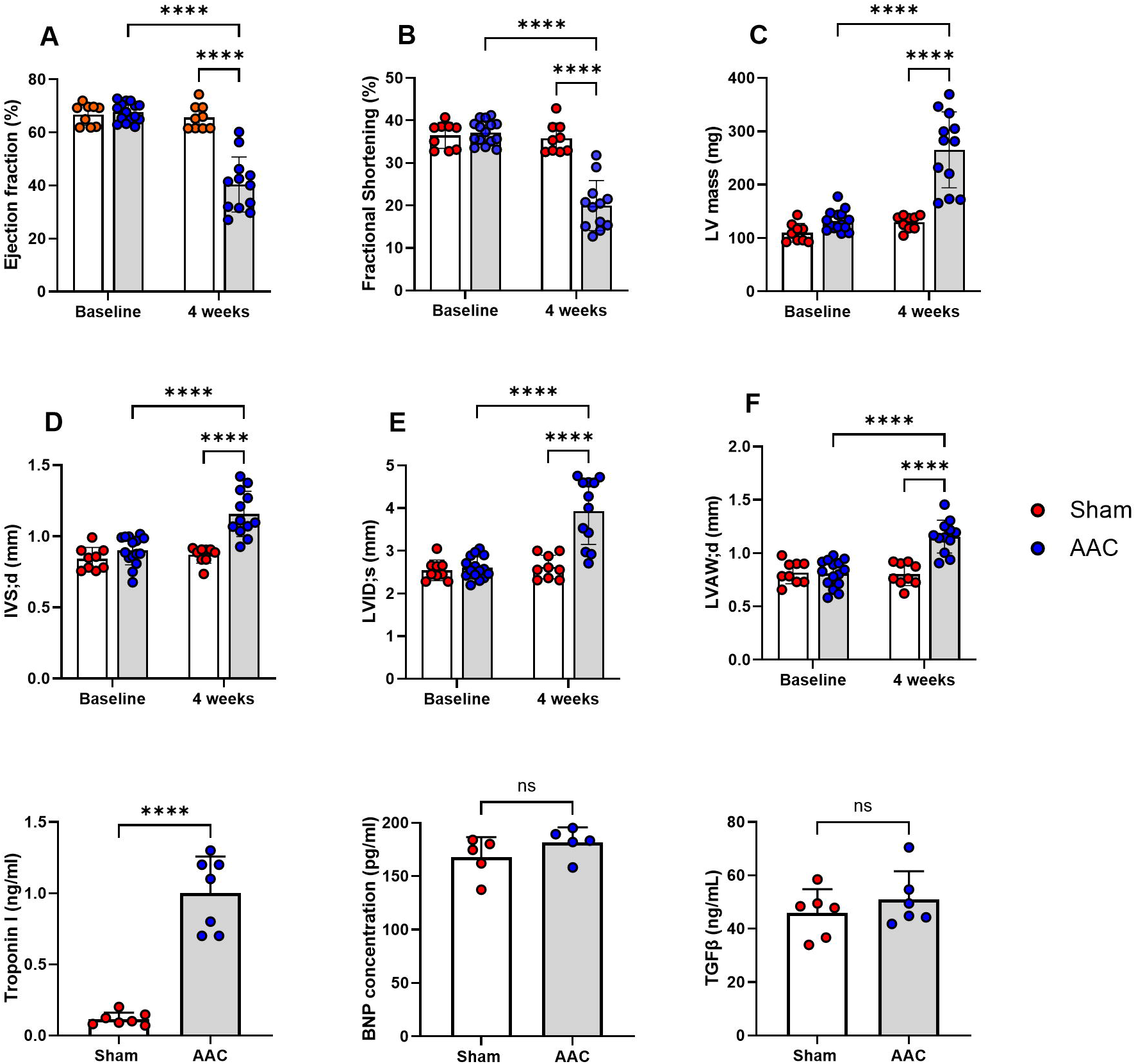
Hemodynamic response measured by echocardiography, 4 weeks after abdominal aortic constriction. IVS;d & LVID;s=left ventricular internal diameter in systole and diastole respectively, LVAW;d = left ventricular anterior wall thickness in diastole. Bottom row: Blood biomarkers of cardiac injury: troponin I, brain natriuretic peptide, tumour growth factor β.

**Figure 2.**
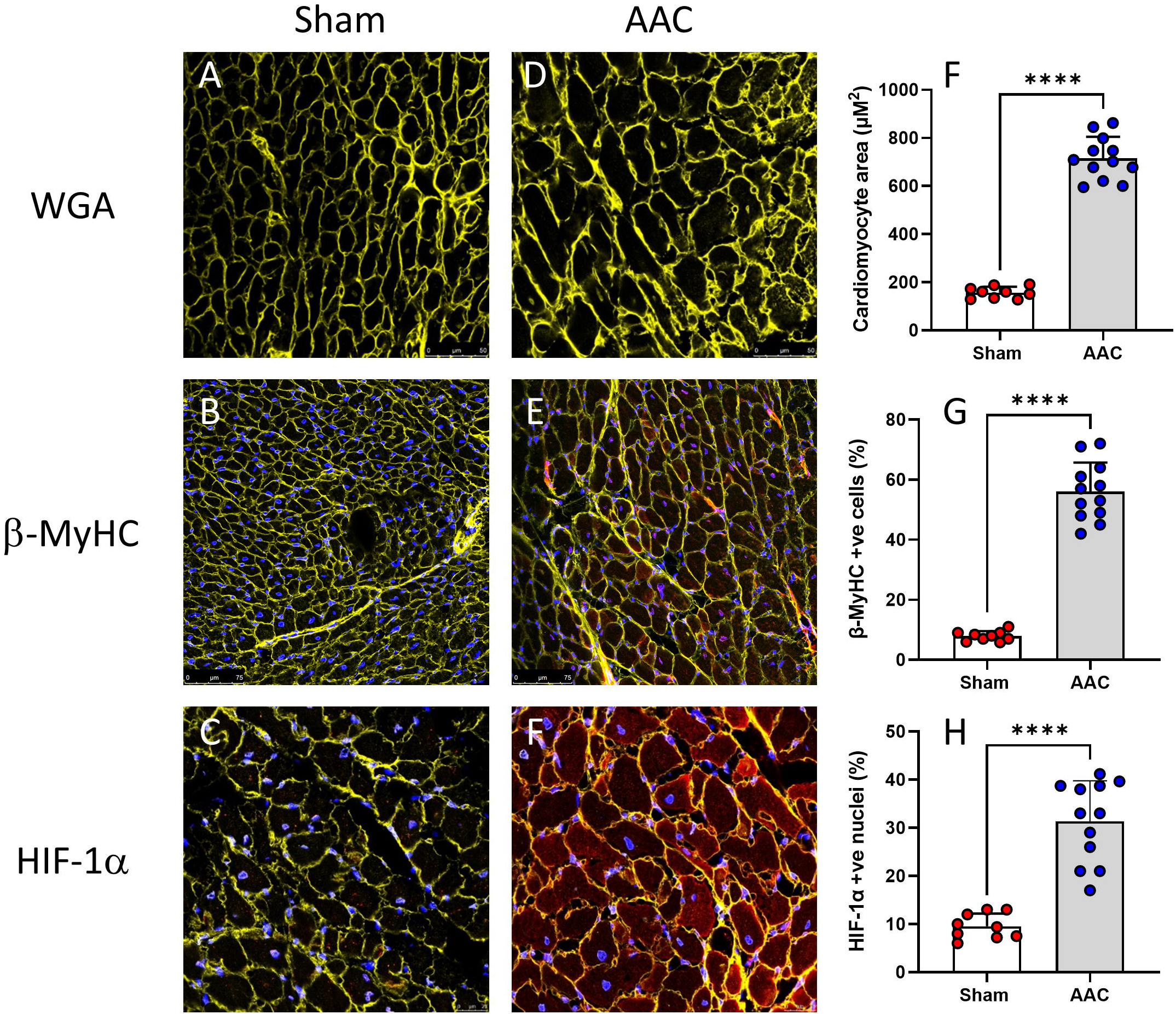
Hypertrophic response by histology, 4 weeks after abdominal aortic constriction. Representative images of histology and summarized quantification from hearts from AAC-operated animals (grey bars, blue datapoints), with time-matched sham controls (open bars, red datapoints). Rows from top to bottom: cardiomyocyte size delineated by wheat germ agglutinin (WGA), β-myosin heavy chain expression (β-MyHC) and nuclear hypoxia-inducible factor 1α (HIF1α) staining (mean ± SD, unpaired t-test, *p<0.05, **p<0.001, ****p<0.0001).

### Evaluation of [^64^Cu]CuCTS PET for imaging hypoxia in AAC-induced cardiac hypertrophy

Cardiac hypertrophy led to significantly elevated cardiac deposition of ^64^Cu from [^64^Cu]CuCTS from 0.48 ± 0.22 to 1.47 ± 0.93 by PET imaging (p = 0.013) and 1.76 ± 0.33 to 3.13 ± 0.71 (p>0.001) by ex vivo organ counting (Figure 3), despite there being no corresponding change in the cardiac uptake of the perfusion tracer [^64^Cu]CuGTSM by either imaging (35.38 ± 4.78 to 39.56 ± 4.23) or ex vivo counting (42.98 ± 4.77 to 46.39 ± 6.69). AAC surgery resulted in a broad range of hypertrophic responses, but cardiac [^64^Cu]CuCTS uptake correlated strongly on an animal-by-animal basis with degree of cardiac contractile dysfunction (by ejection fraction R^2^ = 0.513, p=0.008), cardiac hypertrophy (by heart weight: body weight ratio R^2^ = 0.458, p = 0.002) and HIF1α stabilisation (R^2^ = 0.761, p<0.0001) (Fig 4).

**Figure 3.**
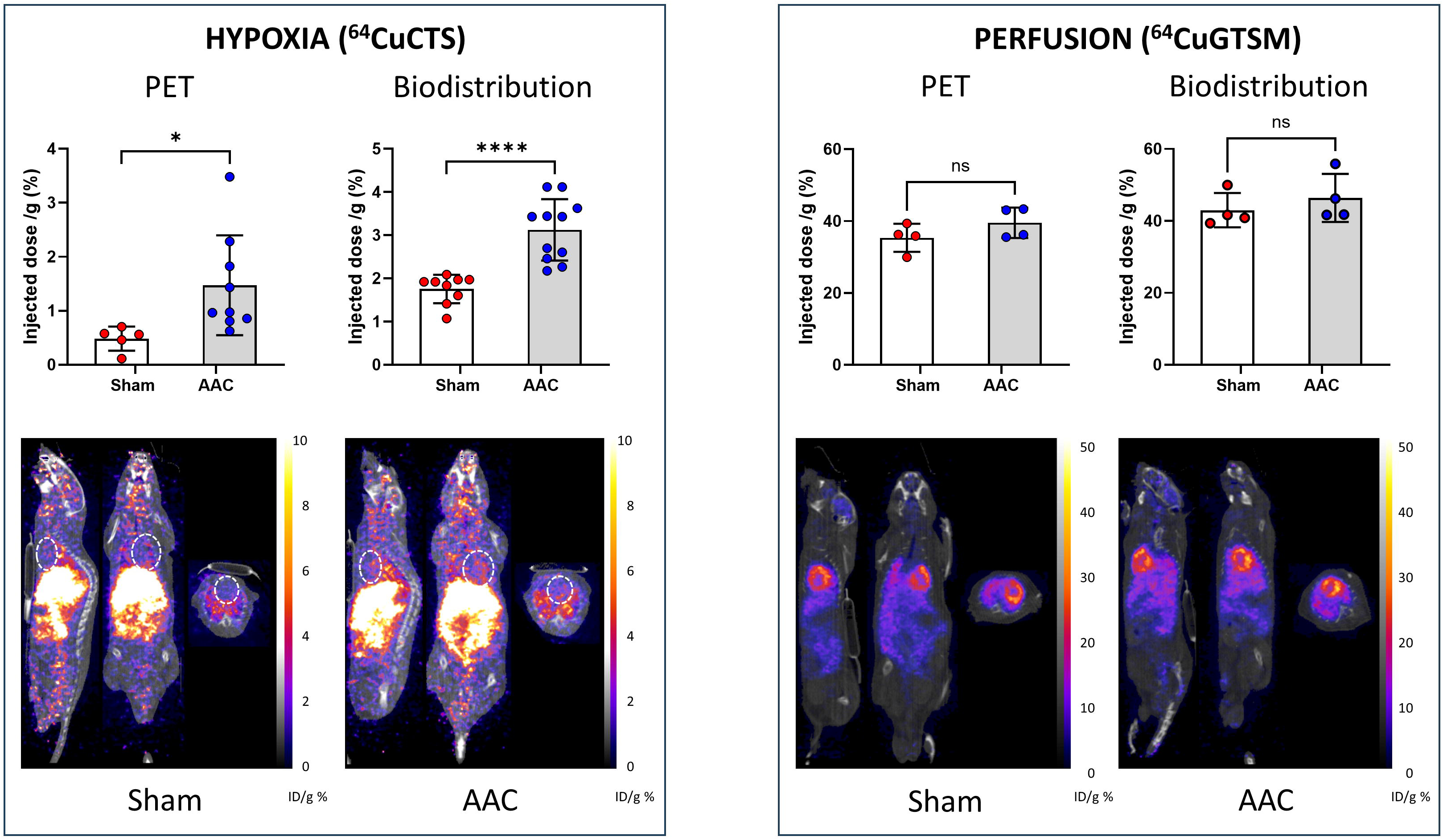
PET imaging of hypoxia and perfusion (with corresponding biodistribution data) in hypertrophic myocardium. Top left: Representative PET images obtained using the hypoxia tracer [^64^Cu]CuCTS 4 weeks after AAC surgery (grey bars, blue datapoints), with corresponding shams (open bars, red datapoints). Graphs show cardiac tracer retention from imaging data (left) and gamma counting of excised hearts (right). Top right: corresponding data obtained with the perfusion tracer [^64^Cu]CuGTSM. Datapoints from AAC-instrumented animals are blue, sham animal datapoints are red (mean ± SD, unpaired t-test except two-way ANOVA analysis with Tukey correction for g, h; *p<0.05, **p<0.001, ****p<0.0001).

**Figure 4.**
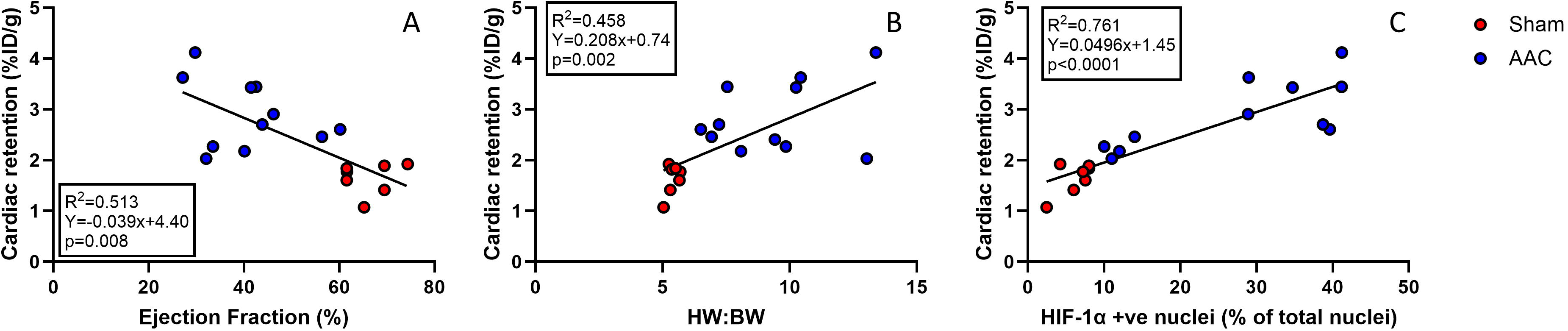
Correlation between cardiac [^64^Cu]CuCTS retention by post-mortem γ-counting with cardiac ejection fraction (left), heart weight:body weight ratio (centre) and nuclear HIF1α staining (right). Datapoints from individual animals 4 weeks post AAC or sham are shown in blue and red respectively.

Whole body [^64^Cu]CuCTS and [^64^Cu]CuGTSM biodistributions are displayed in the **data supplement**. ^64^Cu deposition from CTS was lower in the small intestine in AAC-operated animals, but no other observable non-cardiac differences were noted between treatment groups.

### Dysregulated metabolism in AAC-induced hypertrophy

Hypertrophic myocardium exhibited significant metabolic derangement, characterized by depleted phosphocreatine, ATP and fumarate and elevated tissue succinate and lactate (full details in **data supplement table 1**). Unsupervised hierarchical clustering associated [^64^Cu]CuCTS accumulation (by both imaging and ex vivo counting) most strongly with elevated cardiac succinate and lactate and depleted ATP, acetyl carnitine, fumarate and NAD^+^ (Fig 5). The analysis resolved the animals that had undergone ACC from the sham animals, except for three individuals which clustered with the sham group despite having undergone AAC surgery. These mice exhibited no metabolic perturbation, no stabilisation of cardiac HIF1α, and no elevated cardiac [^64^Cu]CuCTS accumulation. The lack of a hypertrophic response in these animals was mirrored in the principal component analysis of the cohort, where these animals more closely mimicked the sham cohort, particularly with respect to succinate, lactate and fumarate profiles and low [^64^Cu]CuCTS retention. This highlights the inherent variability of the model, and the ability of [^64^Cu]CuCTS to distinguish hypoxic from normal myocardium, even in the AAC-operated cohort.

**Figure 5.**
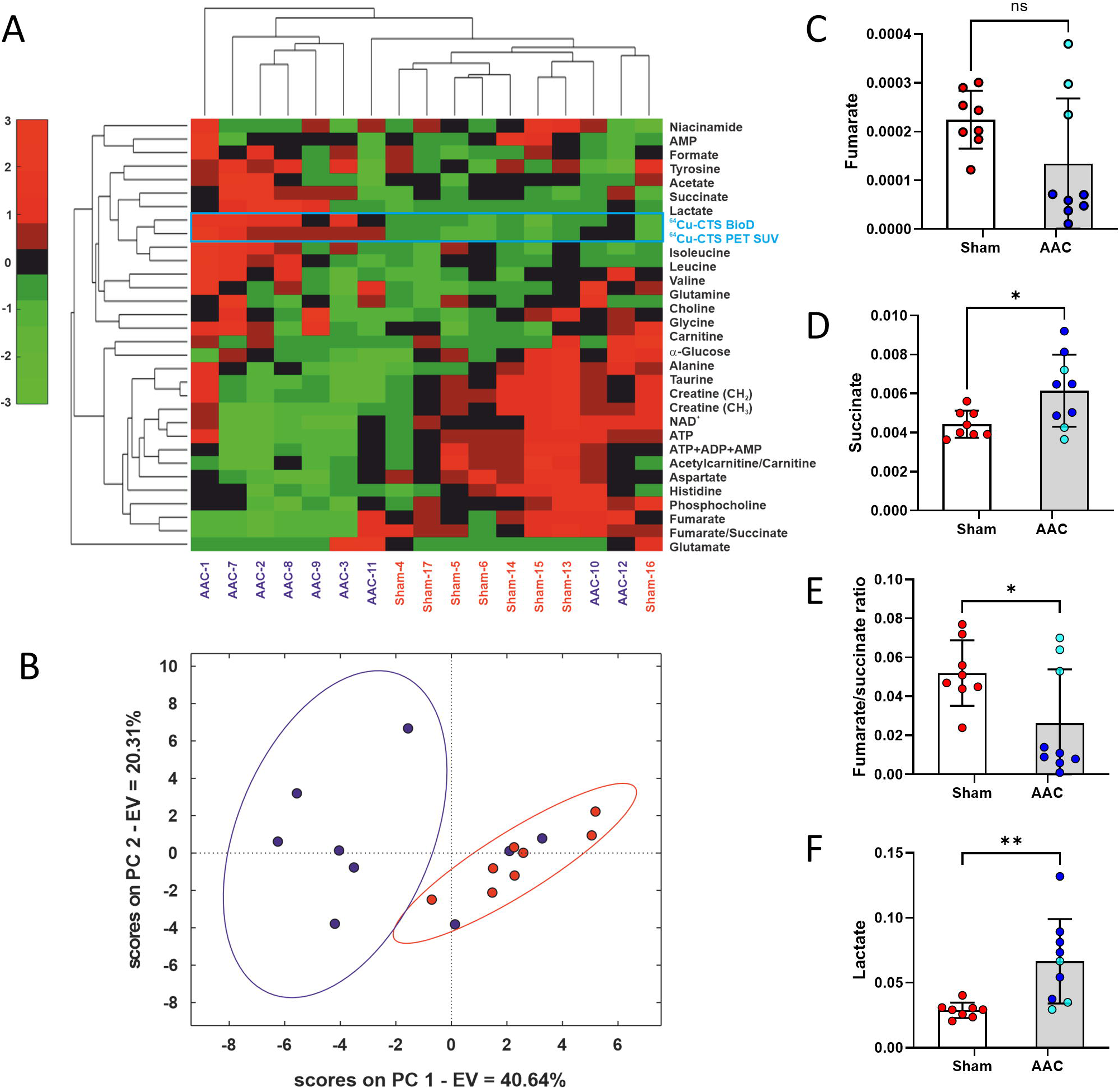
Correlation between cardiac [^64^Cu]CuCTS retention by post-mortem γ-counting and metabolomic data. Top left: Hierarchical cluster analysis showing animal-by-animal association between cardiac metabolomic data, [^64^Cu]CuCTS retention (y-axis) and animal treatment. Individual animals on the x-axis are either 4 weeks post AAC-surgery or time-matched shams. Bottom left: PCA showing 3 of the AAC animals (blue) clustering with the shams (red). Right: Graphs showing key metabolic changes (fumarate, succinate & lactate) in AAC-induced hypertrophy. The 3 AAC animals which clustered with the shams are highlighted with sky blue datapoints.

**Figure 6.**
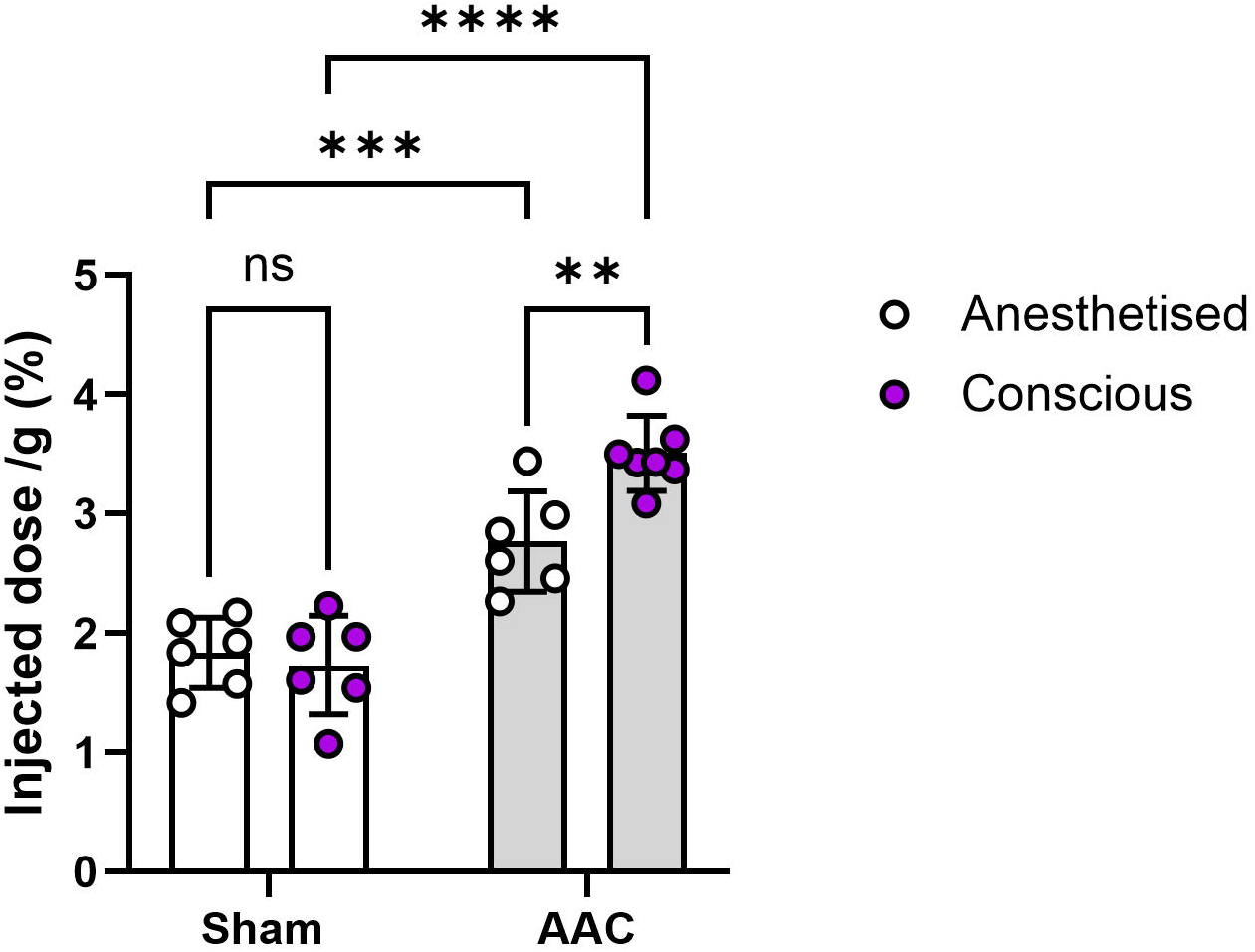
The effect of anesthesia on cardiac hypoxia as measured by [^64^Cu]CuCTS retention. 4 weeks after ACC surgery (grey bars) or sham surgery (open bars), mice were either injected with [^64^Cu]CuCTS whilst conscious using a restrainer (pink datapoints), or anesthetized with isofluorane (white datapoints). 20 minutes later, all animals were anesthetized for quantification of cardiac radiotracer retention by PET and post-mortem radiotracer biodistribution.

### The effect of anesthesia on cardiac retention of [^64^Cu]CuCTS

In AAC mice, cardiac [^64^Cu]CuCTS uptake was significantly higher in conscious mice than in anesthetised mice (3.49 ± 0.38 vs 2.77 ± 0.42 ID/g (p < 0.0001), while in sham mice cardiac [^64^Cu]CuCTS uptake was not significantly different (1.84 ± 0.29 vs 1.86 ± 0.29) (Figure 5).

## Discussion

We have demonstrated that the PET radiotracer [^64^Cu]CuCTS can non-invasively quantify hypoxia in hypertrophic myocardium *in vivo.* The imaging readout that it provides correlates on an animal-by-animal basis with the degree of cardiac hypertrophy and contractile dysfunction, the severity of metabolic derangement and the stabilisation of HIF1α. Using [^64^Cu]CuGTSM PET perfusion imaging, we demonstrate that this hypoxia is at the cellular level, and not caused by measurably impaired perfusion. We therefore suggest that [^64^Cu]CuCTS offers promising new opportunities for the detection and stratification of cardiovascular disease using hypoxia-targeting PET imaging.

Bisthiosemicarbazone (BTSC) PET imaging relies on the redox-dependent trapping of radiocopper. As small lipophilic radiocopper complexes, Cu(II)BTSCs are highly cell penetrant, and once inside a cell they are liable to reduction to unstable Cu(I) complexes. When oxygen is abundant they are rapidly re-oxidised to their native Cu(II) form, which is able to leave the cell(21). If tissue oxygen is insufficient, the Cu(I) species dissociate, depositing their radiocopper payload inside the cell in a hypoxia-dependent manner, where it becomes sequestered by copper chaperone proteins(22).

A useful feature of these complexes is their tuneability. By modifying their redox potential they can be made sensitive to different degrees of hypoxia, while modifying their lipophilicity alters their pharmacokinetic properties and cell penetration(23). [^64^Cu]CuATSM is the best known of this library of complexes, but we and others have shown that it is better suited to the extreme hypoxia associated with the cancers that it was designed to detect(24), rather than the more subtle hypoxia driving chronic cardiovascular disease (25). It is interesting that the redox potentials of [^64^Cu]CuCTS and [^64^Cu]CuATSM are the same (−0.59V vs. Ag/AgCl), but [^64^Cu]CuCTS appears more suitable for detecting clinically relevant degrees of hypoxia than its predecessor [^64^Cu]CuATSM. We suggest that its lower lipophilicity (logP 1.31 vs 1.69 in octanol/water) allows it to penetrate deeper into hypoxic tissues, with less non-selective accumulation in cell membranes, to deliver greater hypoxia sensitivity and hypoxic:normoxic contrast (13). By comparison, [^64^Cu]CuPTSM, with a redox potential of −0.51V, deposits radiocopper even in normoxic tissues. Density Functional Theory calculations and electrochemical experiments suggest that less alkylated complexes like CuPTSM and CuGTSM may dissociate more rapidly (precluding the opportunity for oxygen “sensing”), whereas the doubly-alkylated backbones of complexes like CuATSM and CuCTS allow the reversible formation of intermediate Cu(I) complexes which can be re-oxidised by oxygen to provide hypoxia-dependent contrast (26). Thus, while the intracellular reductants for these complexes are shared, it is their relative susceptibility to re-oxidation which governs the hypoxia-dependent imaging contrast that each delivers. While [^64^Cu]CuPTSM has long been proposed as a cardiac perfusion imaging agent (27), more recent studies show that it has a slight hypoxia-selective component to its cellular uptake (24). We have therefore used [^64^Cu]CuGTSM because with a redox potential of −0.43V it dissociates even more rapidly to quantify perfusion with high extraction fraction and least sensitivity to hypoxia (28).

Although we cannot use our approach to quantify absolute intracellular oxygen concentration, we would argue that this parameter would only be useful if the oxygen thresholds at which critical biochemical processes are compromised *in vivo* are also known (which is rarely true). Arguably, the most appropriate technique for detecting prognostically useful energetic compromise in ischemic cardiac syndromes would be ^31^P NMR spectroscopy(29), but it is not widespread because it is technically challenging and its poor sensitivity results in poor spatial and temporal resolution and long scan times. Our approach of correlating [^64^Cu]CuCTS uptake with cardiac energetics could provide similarly useful mechanistic information for identification and sub-stratification of disease but without these limitations. We have previously shown in isolated perfused hearts that [^64^Cu]CuCTS identifies myocardium at the critical threshold for hypoxic compromise, correlating with the onset of anaerobic glycolysis, HIF1α stabilization and loss of contractile performance (17). We have also used ^31^P NMR spectroscopy to show that cardiac [^64^Cu]CuCTS uptake corresponds to the onset of loss of cardiac phosphocreatine and ATP levels, and could thus be considered a proxy for ^31^P NMR spectroscopy but with shorter scan times and higher spatial resolution. We show here that this capability translates *in vivo*. The degree of hypertrophy that we observe in response to AAC surgery is somewhat variable, being dependent on how tightly the suture is tied in each case and how large each mouse was at the time of surgery, which impact on the degree of aortic restriction evoked and the degree of hypertrophy that results. Somewhat ironically, this inherent variability in our model highlights the utility and advantages of [^64^Cu]CuCTS PET imaging. On an animal-by-animal basis, metabolic perturbation (manifesting the classic ischemic/hypoxic phenotype of mitochondrial compromise and elevated anaerobic glycolysis) and HIF1α stabilization all correlated with each other as one might expect, with the most hypertrophic hearts displaying the most derangement and vice versa. Unsupervised hierarchical cluster analysis showed that ^64^Cu accumulation delivered by [^64^Cu]CuCTS correlated with these parameters. Moreover, in the animals which underwent AAC surgery but did not hypertrophy (presumably because the ligature wasn’t sufficiently tight), there was no metabolic perturbation, no HIF1α stabilization and no elevated [^64^Cu]CuCTS uptake. As such, [^64^Cu]CuCTS PET can highlight the *pathophysiological relevance* of a flow restriction (i.e. in this case the induction of hypoxia), rather than just identifying the restriction itself.

## Study Limitations

We have radiolabeled our ligands with ^64^Cu in this study because of its relatively long half-life, which has advantages practically and economically for preclinical experimental work. While its uptake can be accurately quantified by *ex vivo* biodistribution for proof-of-concept as we have done, ^64^Cu a low positron yield (19%) and a relatively long half-life (12.7 hours), which mean that dosimetry is not ideal, and its β^+^ energy (657 KeV) is not ideal for preclinical work where its relatively poor resolution makes imaging mouse hearts challenging, as our images demonstrate. However, this would be less of a limitation in human hearts in clinical scanners, and these tracers rapidly clear from blood to provide images with adequate target-to-background ratio within 15 minutes, which make them amenable for radiolabelling with ^62^Cu instead. With a 97% positron yield and 9.7 minutes half-life, this would result in a significantly lower dose to the patient (30). We will investigate this possibility, and the potential of [^64^Cu/^62^Cu]CuCTS to detect cardiac hypoxia *<u>before</u>* contractile dysfunction manifests, in future work.

## Clinical Perspectives

There is currently much focus on the optimization of imaging techniques for the characterization of cardiovascular disease in terms of perfusion or perfusion reserve (31,32). While useful for characterizing coronary artery disease, these approaches provide little reference to the energetic or oxygen demands of subtended tissues or the pathophysiological importance of any measured perfusion deficit. Similarly, while BOLD MRI can measure cardiac blood oxygenation, this is hard to relate to oxygen saturation in cardiac myocytes themselves to determine whether cardiac oxygen *demand* is being met. As such, these techniques may be sub-optimal for sub-stratifying patients with less severe coronary artery occlusions, or microvascular dysfunction associated with angina, INOCA (33), HFpEF, pulmonary artery hypertension (34) or cardiotoxicities associated with cancer therapies (35), where ischemia is subtle and diffuse. This distinction is increasingly being appreciated, with recent studies re-evaluating the relevance of exercise stress testing to reveal ischemic myocardium in the absence of measurable perfusion deficit (36). Our data suggest that these low-grade chronic ischemic syndromes could be more sensitively and specifically characterized using molecular imaging techniques like that proposed here rather than the functional imaging techniques currently used.

## Abbreviations

AAC: abdominal aortic constriction
BTSC: bisthiosemicarbazone
EF: ejection fraction
FAC: fractional area change
FS: fractional shortening
HIF: hypoxia inducible factor
IVS: intraventricular septum
LVID: left ventricular internal diameter
LVAW: left ventricular anterior wall
LVPW: left ventricular posterior wall
PET: positron emission tomography

---

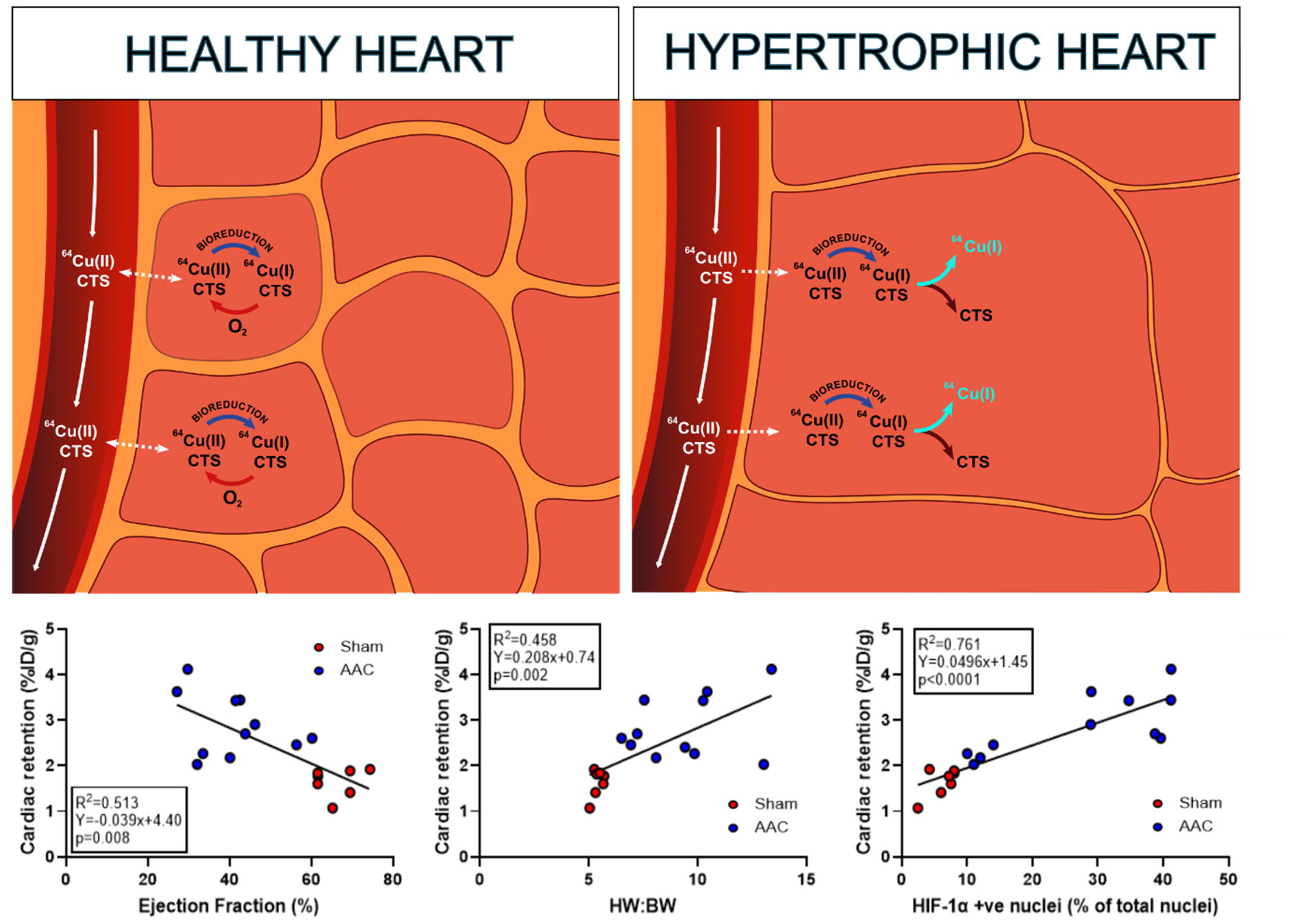

## REFERENCES

1. Lanza GA, Crea F. Primary Coronary Microvascular Dysfunction. Circulation 2010;121:2317–2325.

2. Giordano FJ. Oxygen, oxidative stress, hypoxia, and heart failure. J. Clin. Invest. 2005;115:500–8.

3. Liu Y, Cox SR, Morita T, Kourembanas S. Hypoxia Regulates Vascular Endothelial Growth Factor Gene Expression in Endothelial Cells. Circ. Res. 1995;77:638–643.

4. Semenza GL. Hypoxia-inducible factors: roles in cardiovascular disease progression, prevention, and treatment. Cardiovasc. Res. 2022;119:371–380.

5. Bishop T, Ratcliffe PJ. HIF hydroxylase pathways in cardiovascular physiology and medicine. Circ. Res. 2015;117:65–79.

6. Takahashi N, Fujibayashi Y, Yonekura Y et al. Copper-62 ATSM as a hypoxic tissue tracer in myocardial ischemia. Annals Nucl. Med. 2001;15:293–6.

7. Hearse DJ. Myocardial ischaemia: can we agree on a definition for the 21st century? Cardiovasc. Res. 1994;28:1737–1744.

8. Serio S, Pagiatakis C, Musolino E et al. Cardiac Aging Is Promoted by Pseudohypoxia Increasing p300-Induced Glycolysis. Circ. Res. 2023;133:687–703.

9. Marfella R, D’Amico M, Di Filippo C et al. Myocardial infarction in diabetic rats: role of hyperglycaemia on infarct size and early expression of hypoxia-inducible factor 1. Diabetologia 2002;45:1172–1181.

10. Schindler TH, Fearon WF, Pelletier-Galarneau M et al. Myocardial Perfusion PET for the Detection and Reporting of Coronary Microvascular Dysfunction: A JACC: Cardiovascular Imaging Expert Panel Statement. J Am Coll Cardiol Img. 2023;16:536–548.

11. Sinusas AJ. The potential for myocardial imaging with hypoxia markers. Sem. Nucl. Med. 1999;29:330–8.

12. Xie F, Wei W. [(64)Cu]Cu-ATSM: an emerging theranostic agent for cancer and neuroinflammation. Eur. J. Nucl. Med. Mol. Imaging 2022;49:3964–3972.

13. Handley MG, Medina RA, Paul RL, et al. Demonstration of the retention of 64Cu-ATSM in cardiac myocytes using a novel incubation chamber for screening hypoxia-dependent radiotracers. Nucl. Med. Comms 2013;34:1015–22.

14. Handley MG, Medina RA, Mariotti E et al. Cardiac hypoxia imaging: second generation analogues of 64Cu-ATSM. J. Nucl. Med. 2014;55:488–494.

15. Lewis JS, Herrero P, Sharp TL et al. Delineation of hypoxia in canine myocardium using PET and copper(II)-diacetyl-bis(N4-methylthiosemicarbazone). J. Nucl. Med. 2002;43:1557–1569.

16. Takahashi N, Fujibayashi Y, Yonekura Y et al. Copper-62 ATSM as a hypoxic tissue tracer in myocardial ischemia. Ann. Nucl. Med. 2001;15:293–6.

17. Medina RA, Mariotti E, Pavlovic D et al. 64Cu-CTS: A Promising Radiopharmaceutical for the Identification of Low-Grade Cardiac Hypoxia by PET. J. Nucl. Med. 2015;56:921–926.

18. Jauregui-Osoro M, De Robertis S, Halsted P et al. Production of copper-64 using a hospital cyclotron: targetry, purification and quality analysis. Nucl. Med. Comms. 2021;42:1024–1038.

19. Loonat AA, Martin ED, Sarafraz-Shekary N et al. p38γ MAPK contributes to left ventricular remodeling after pathologic stress and disinhibits calpain through phosphorylation of calpastatin. FASEB journal. 2019;33:13131–13144.

20. Beckonert O, Keun HC, Ebbels TM et al. Metabolic profiling, metabolomic and metabonomic procedures for NMR spectroscopy of urine, plasma, serum and tissue extracts. Nature protocols 2007;2:2692–703.

21. Dearling JLJ, Lewis JS, Mullen GED et al. Design of hypoxia-targeting radiopharmaceuticals: selective uptake of copper-64 complexes in hypoxic cells in vitro. eJNMMI Res 1998;25:788–792.

22. Dearling JLJ, Packard AB. On the Destiny of (Copper) Species. J Nucl Med 2014;55:7–8.

23. Blower PJ, Castle TC, Cowley AR et al. Structural trends in copper(ii) bis(thiosemicarbazone) radiopharmaceuticals. Dalton Trans. 2003:4416–4425.

24. Dearling J, Lewis J, Mullen G et.al Copper bis(thiosemicarbazone) complexes as hypoxia imaging agents: structure-activity relationships. J Biol Inorg Chem 2002;7:249–259.

25. Pell VR, Baark F, Mota F et al. PET Imaging of Cardiac Hypoxia: Hitting Hypoxia Where It Hurts. Current Cardiovasc. Imaging Rep. 2018;11:7.

26. Maurer RI, Blower PJ, Dilworth JR et al. Studies on the mechanism of hypoxic selectivity in copper bis(thiosemicarbazone) radiopharmaceuticals. J Med Chem 2002;45:1420–31.

27. Torres JB, Andreozzi EM, Dunn JT et al. PET Imaging of Copper Trafficking in a Mouse Model of Alzheimer Disease. J. Nucl. Med. 2016;57:109–14.

28. Andreozzi EM, Torres JB, Sunassee K et al. Studies of copper trafficking in a mouse model of Alzheimer’s disease by positron emission tomography: comparison of (64)Cu acetate and (64)CuGTSM. Metallomics 2017;9:1622–1633.

29. de Wit-Verheggen VHW, Schrauwen-Hinderling VB, Brouwers K et al. PCr/ATP ratios and mitochondrial function in the heart. A comparative study in humans. Sci Rep 2023;13:8346.

30. Hao G, Singh AN, Oz OK, Sun X. Recent advances in copper radiopharmaceuticals. Current radiopharm. 2011;4:109–21.

31. van Diemen PA, de Winter RW, Schumacher SP et al. The diagnostic performance of quantitative flow ratio and perfusion imaging in patients with prior coronary artery disease. Eur Heart J Cardiovasc. Imaging 2023;25:116–126.

32. Benenati S, Montorfano M, Pica S et al. Coronary physiology thresholds associated with microvascular obstruction in myocardial infarction. Heart 2024;110:271–280.

33. Kaski JC, Crea F, Gersh BJ, Camici PG. Reappraisal of Ischemic Heart Disease. Circulation 2018;138:1463–1480.

34. Johnson S, Sommer N, Cox-Flaherty K et al. Pulmonary ypertension: A Contemporary Review. Am. J Resp. Crit. Care Med. 2023;208:528–548.

35. Baik AH. Hypoxia signaling and oxygen metabolism in cardio-oncology. J. cardiol 2022;165:64–75.

36. Sinha A, Dutta U, Demir OM et al. Rethinking False Positive Exercise Electrocardiographic Stress Tests by Assessing Coronary Microvascular Function. J Am Coll Cardiol 2024;83:291–299.

