## Supplemental methods and data for "Perfusion-Independent Tissue Hypoxia in Cardiac Hypertrophy in Mice Measured by ^64^Cu-CTS PET Imaging"

**Immunohistochemistry**

20 µM sections were cut at -20 °C using a Leica CM 1800 cryostat, and transferred to Superfrost Plus™ Adhesion slides (ThermoFisher). Frozen slides were air dried and fixed with acetone before being rehydrated with PBS-t for 5 mins. Sections were then blocked using 5% BSA for 30 mins before primary antibody (1:200 dilution) was added overnight at 4 °C. Slides were then washed with PBS-t for 5 mins 3 times prior to secondary antibody incubation for 1 hr at room temperature. After a further wash, slides were mounted with prolong gold mounting solution (H-1000, Vector Laboratories). Sections were imaged using a Leica TCS SP5 II, confocal microscope and analysed using LAS X Life Science Software.

**Western blotting**

30 mg samples were added to cold Matrix tubes containing 1.4 mm ceramic beads and RIPA buffer containing 1 x protease and phosphatase inhibitors. Samples were lysed by rapid shaking at 4 °C (Precellys 24 homogeniser, Bertin Instruments), centrifuged at 15,000 x *g* at 4 °C for 10 min, and the supernatant collected. Protein content was determined using a BCA assay kit. NuPAGE™ LDS Sample Buffer and NuPAGE™ reducing agent were added to 80 µg cell lysate in a total volume of 20 µL. Samples were then vortexed and heated to 95 °C for 5 min prior to loading onto 10% polyacrylamide gels alongside a protein ladder (Precision Plus Protein Dual Colour Standards (BioRad) and samples were run at 120 V. Proteins were transferred to PVDF membrane using a Trans-Blot Turbo Transfer System (Bio-Rad). Blots were probed for HIF-1α (1:500; Novus Biologicals) with β -actin as a loading control with HRP-linked anti-rabbit IgG secondary antibody (1:10,000; Abcam). Membranes were washed in 50 mL tris-buffered saline with 0.1% Tween 20 (TBST) for 10 min, 5 times on a shaker. To visualise proteins, 4 mL Amersham™ ECL Prime Western Blotting Detection Reagent (BioRad) was added to each membrane in the dark for 1 min and exposed on CL-XPosure photographic films (Thermo Scientific). Densitometry was performed using Quantity One Software (BioRad), normalised to β-actin.

**[^64^Cu] Biodistribution analysis**

An 500µl aliquot of blood was obtained via cardiac puncture prior to the heart being excised, weighed, and cut mid-ventricle. The apex and lower ventricles were snap-frozen in liquid nitrogen for western blotting and the upper ventricles were embedded in OCT for sectioning and immunohistochemistry. The other major organs, bone and skeletal muscle were harvested, weighed, and counted with a gamma counter (Compugamma, LKB Wallac, Australia), along with standards prepared from injected material.

**NMR acquisition**

^1^H nuclear magnetic resonance spectra were acquired using a vertical-bore, ultra-shielded Bruker 14.1 tesla (600 MHz) spectrometer equipped with a prodigy probe, at 298K using the Bruker noesygppr1d pulse sequence for residual water suppression. Acquisition parameters were: 64 transients; 4 dummy transients; 20.8 ppm spectral width; acquisition time, 2.6 s; pre-scan delay, 4 s; 90° flip angle; and experiment duration of 7.5 min per sample. TopSpin (version 4.0.5) software was used for data acquisition and for metabolite quantification. Free induction decays (FIDs) were multiplied by a line broadening factor of 0.3 Hz and Fourier transformed, phase, and automatic baseline-corrected. Chemical shifts were normalised by setting the TSP signal to 0.0 ppm. Metabolite peaks of interest were initially integrated automatically using a pre-written integration-region text file and then manually adjusted as required. Assignment of peaks to their respective metabolites was based on previously obtained in-house data, confirmed by chemical shift, and using Chenomx NMR Profiler Version 8.1 (Chenomx, Canada). Metabolite concentrations are normalised to total spectrum area.

**SUPPLEMENTAL DATA**

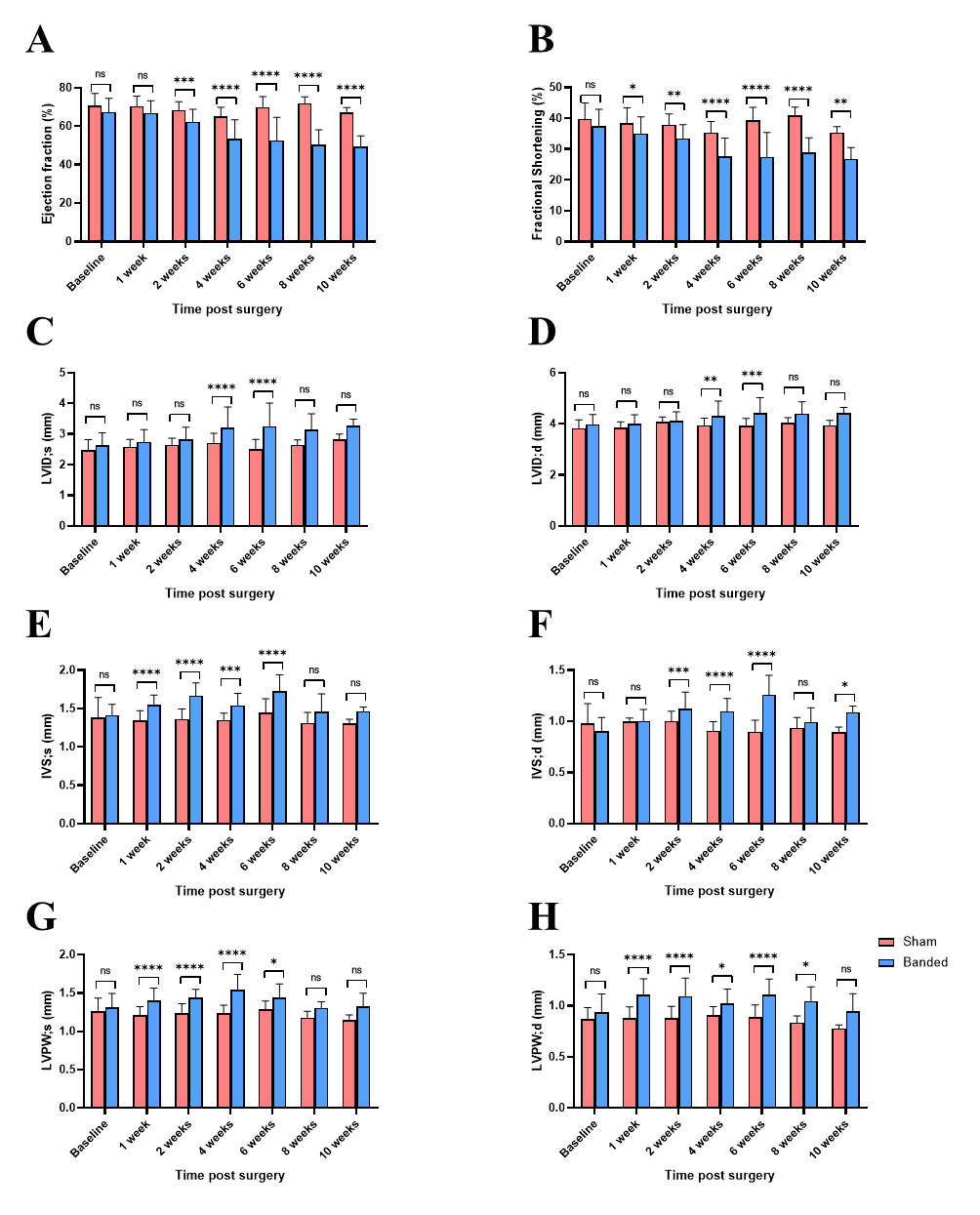

**Supplemental Figure 1:** Longitudinal changes in cardiac hemodynamics following abdominal aortic constriction surgery (pink bars) versus time matched sham controls (blue bars). LVID;s & LVID;d = left ventricular internal diameter at systole & diastole respectively, IVS;s & IVS;s = interventricular wall thickness at systole & diastole respectively, LVPW;s & LVPW;d = left ventricular posterior wall thickness at systole and diastole respectively. n=6/group. *p<0.05, **p<0.01, ***p<0.005, ****p<0.001, ns=not significantly different.

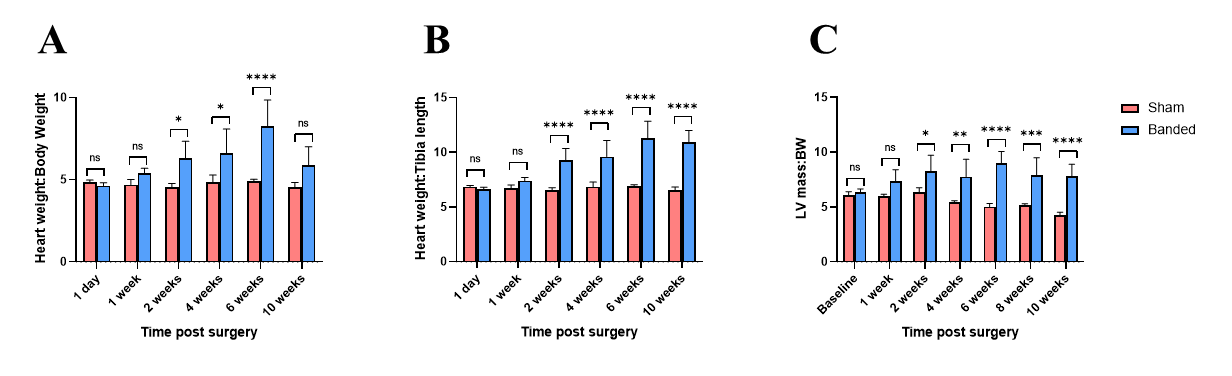

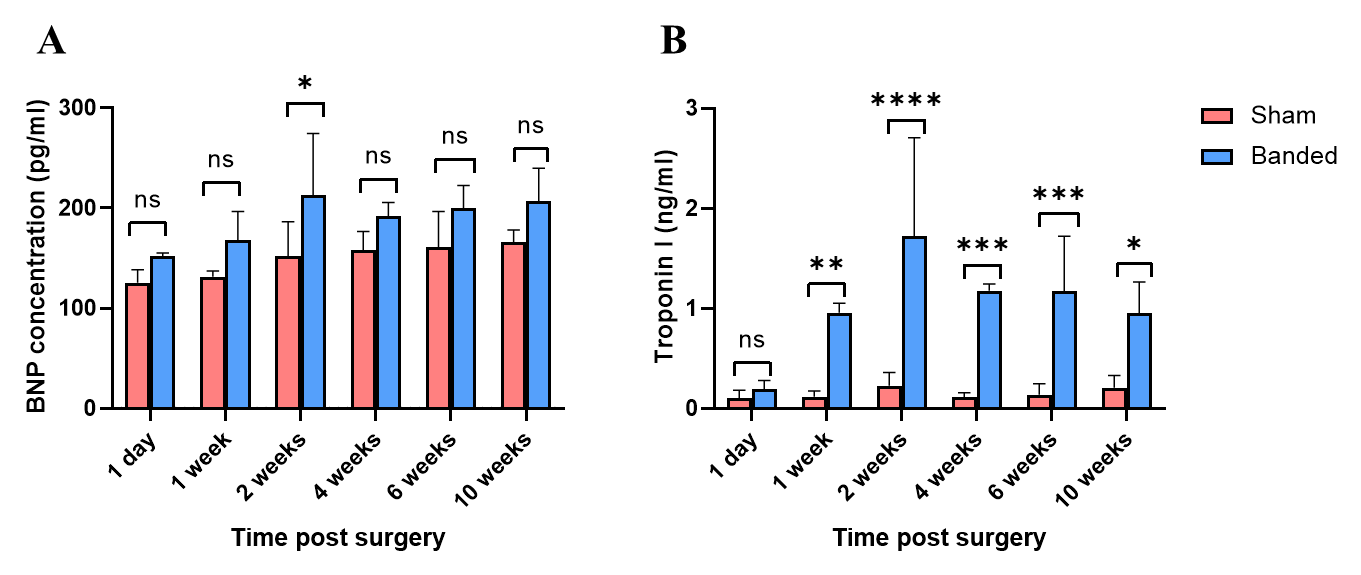

**Supplemental Figure 2:** Longitudinal changes in cardiac hypertrophy and blood biomarkers following abdominal aortic constriction surgery (pink bars) versus time matched sham controls (blue bars). Top graph = changes in heart weight:tibia length, bottom left blood Brain Natruiretic Peptide concentration, bottom right: blood troponin I concentration. n=6/group. *p<0.05, **p<0.01, ***p<0.005, ****p<0.001, ns=not significantly different.

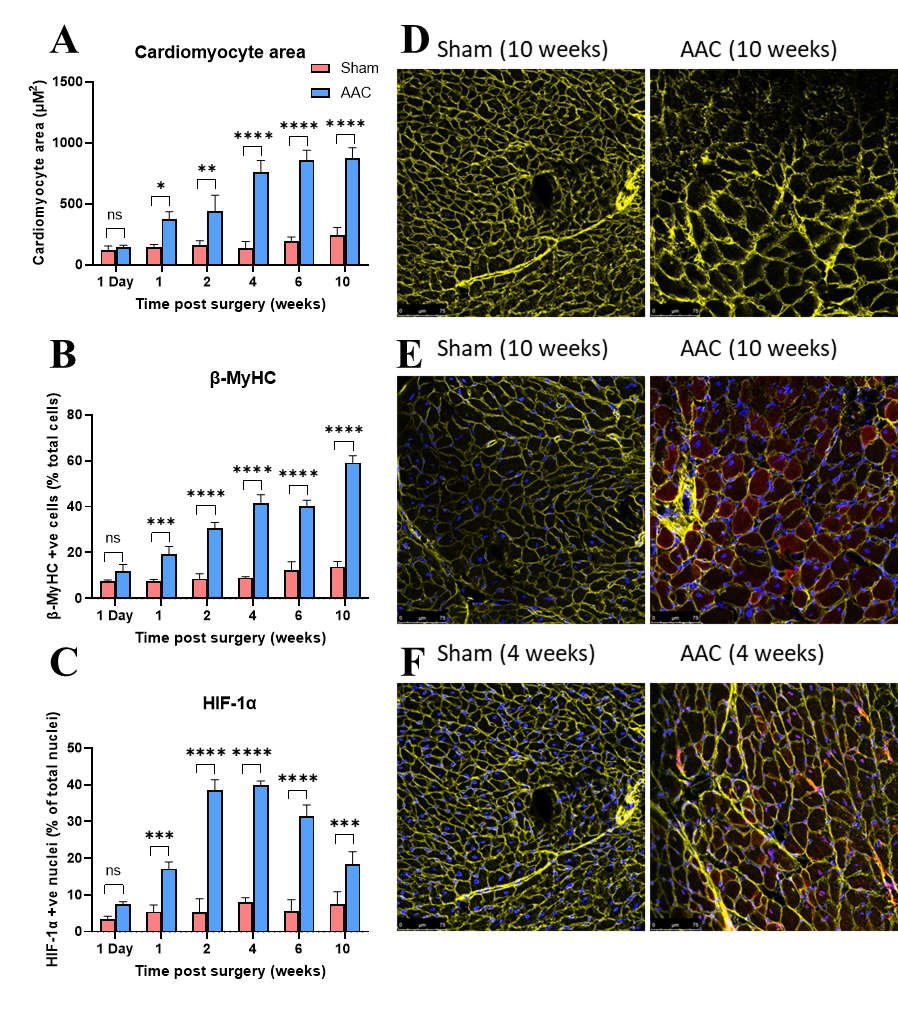
.

**Supplemental Figure 3:** Longitudinal changes in histological biomarkers following abdominal aortic constriction surgery (pink bars) versus time matched sham controls (blue bars). Top row = changes in cardiomyocyte area calculated from wheatgerm agglutinin staining, middle row: b-myosin heavy chain staining, bottom row nuclear HIF1a staining. n=6/group. *p<0.05, **p<0.01, ***p<0.005, ****p<0.001, ns=not significantly different

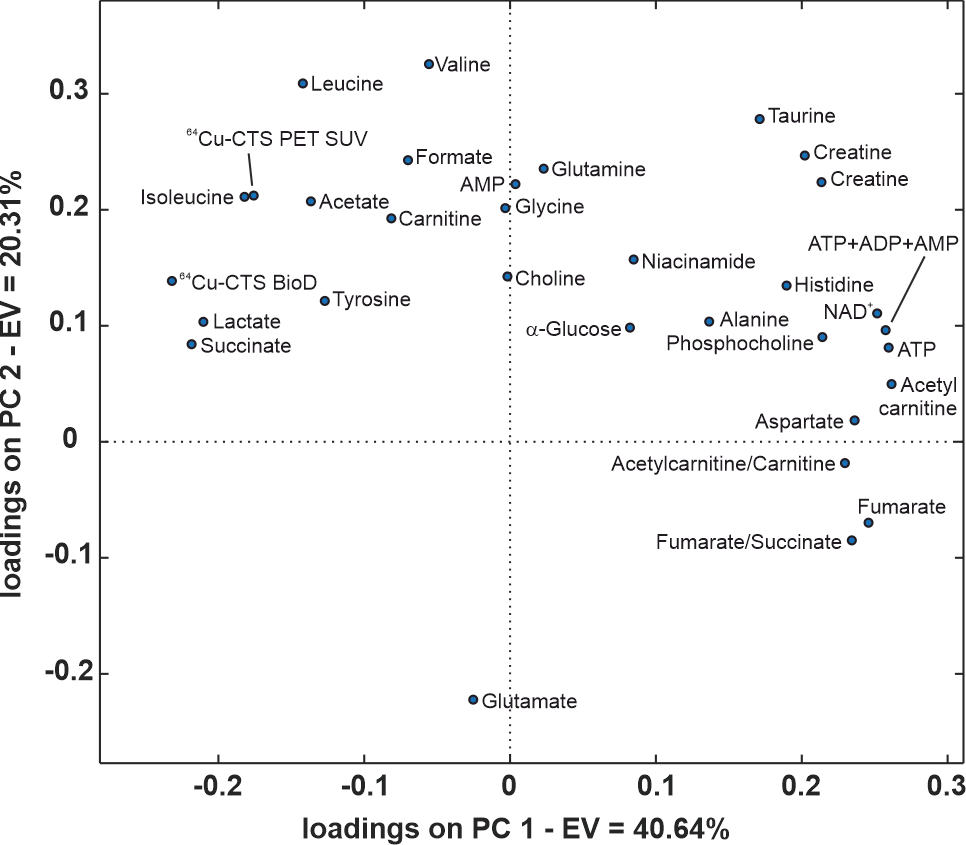

**Supplemental Figure 4:** Principal Component Analysis loadings plot showing the relative contribution of each metabolite/imaging biomarker to the hypoxia-dependent separations displayed in Figure 5.

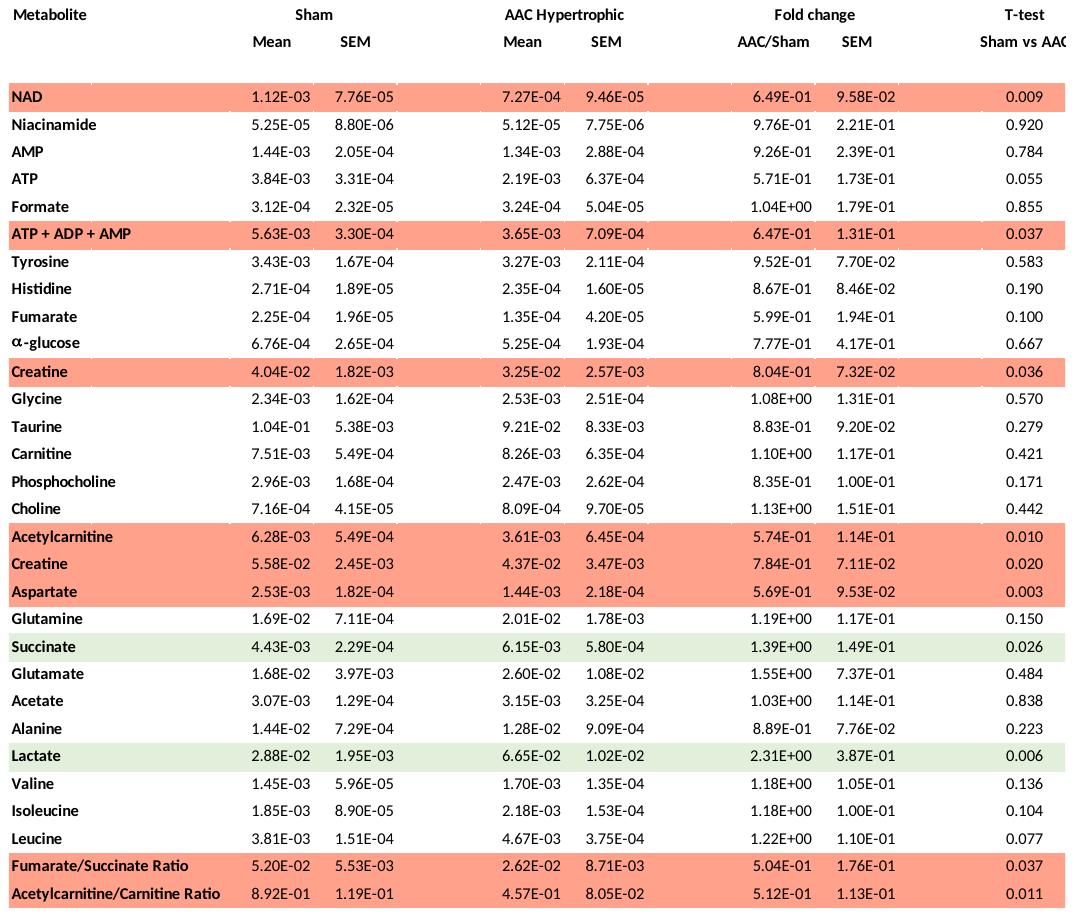

**Supplemental table 1**. Changes in cardiac metabolomic profile 4 weeks after abdominal aortic banding compared with time-matched sham control animals. Rows in red show metabolites significantly lower in AAC (n=9) hearts than sham (n=8), rows in green show metabolites significantly higher in AAC hearts than sham (p<0.05)

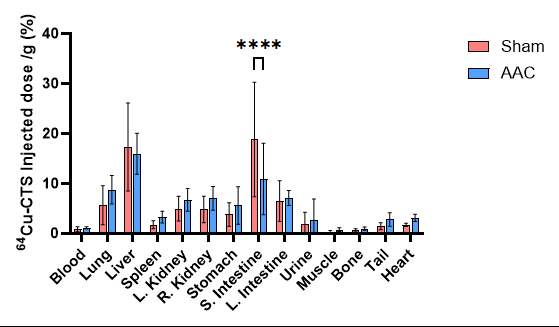

**Supplemental Figure 5.** Whole body post-mortem [^64^Cu]CuCTS biodistributions 4 weeks after abdominal aortic constriction surgery (pink bars) compared to time-matched sham controls (blue bars). n=6/group. ***p<0.005

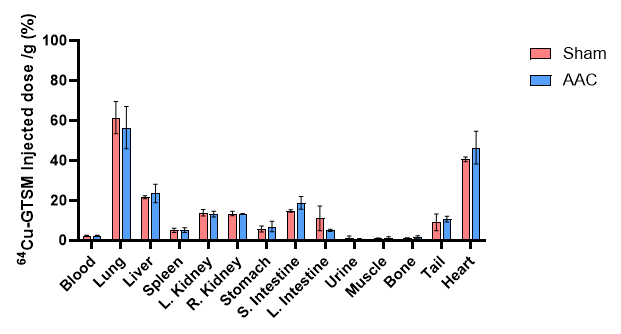

**Supplemental Figure 6.** Whole body post-mortem [^64^Cu]CuGTSM biodistributions 4 weeks after abdominal aortic constriction surgery (pink bars) compared to time-matched sham controls (blue bars). n=6/group.
